# *Arabidopsis* ATHB17 coordinates nuclear and plastidic photosynthesis gene expression in response to abiotic stress

**DOI:** 10.1101/040501

**Authors:** Ping Zhao, Rong Cui, Ping Xu, Jie-Li Mao, Yu Chen, Cong-zhao Zhou, Lin-Hui Yu, Cheng-Bin Xiang

## Abstract

Photosynthesis is sensitive to environmental stresses. How nuclear and plastid genome coordinate to cope with abiotic stress is not well understood. Here we report that ATHB17, an *Arabidopsis* HD-Zip transcription factor, coordinates the expression of nuclear encoded photosynthetic genes (NEPGs) and plastid encoded genes (PEGs) in response to abiotic stress. *ATHB17*-overexpressing plants display enhanced stress tolerance, whereas its knockout mutant is more sensitive compared to the wild type. Through RNA-seq analysis, we found that ATHB17 down-regulated many NEPGs while up-regulated a number of PEGs. ATHB17 could directly modulate the expression of several NEPGs by binding to their promoters. Furthermore, we identified *ATSIG5*, encoding a plastid sigma factor, as one of the target genes of ATHB17. Loss of *ATSIG5* reduced salt tolerance while overexpression of *ATSIG5* enhanced salt tolerance, similar to that of *ATHB17*. Taken together, our results reveal that *ATHB17* is an important coordinator between NEPGs and PEGs partially through ATSIG5 to protect photosynthesis machinery in response to abiotic stresses.

## Introduction

Abiotic stresses such as salinity, drought, high light, unfavorable temperatures, adversely affect the growth and development of plants. Photosynthesis in chloroplast is one of the primary processes to be affected by abiotic stress^1,2^. The effects can be direct, as decreased CO_2_ diffusion caused by stomata close^3^ or affecting ribulose bisphosphate carboxylase/oxygenase (Rubisco) activity^4^. More importantly, abiotic stress reduces the threshold intensity for the onset of photoinhibition, and results in over excitation of the photosystems, thus dramatically increasing reactive oxygen species (ROS) production^5,6^. Rapid response of plant metabolism and photosynthetic machinery is key for plants to cope with the fluctuating environment^7^.

Chloroplasts are genetically semi autonomous organelles that evolutionarily retained eubacteria-type circular genome DNA. In higher plants, the chloroplast 120-150 kb genome encodes only about 120 genes^8^. More required proteins are encoded by the nuclear genome and imported to play roles in the chloroplasts after translation in the cytosol^9^.

Chloroplast gene transcription in higher plants is performed by at least two types of RNA polymerases, plastid encoded RNA polymerase (PEP) and nuclear encoded RNA polymerase (NEP). PEP holoenzyme is a complex composed by five kind of subunits: α, β, β’, ω (omega), σ(sigma), in which the α2ββ’ω constitutes the catalytic core, while the σ subunit recognizes the specific promoter region and initiates transcription to the core complex^10^.

The transcription of photosynthesis related genes in chloroplasts is mainly dependent on PEP, and the nuclear-encoded sigma factors play special roles in regulating the chloroplast transcription^11,12^. Since the first chloroplast sigma factor gene was isolated from red algae nuclear genome^13,14^, more and more chloroplast sigma factors *(ATSIG1-6)* had been identified in different plant species^15,16^. Most plastid encoded genes appear to be regulated by several sigma factors with overlapping functions. However, within a certain time frame during plant development, plastid genes likely to be coordinated by a distinct sigma factor, for example, *PSAA* and *RBCL* by *ATSIG1* ^17^, *PSAJ* by *ATSIG2^18^*, *PSBN* by *ATSIG3^19^*, *NDHF* by *ATSIG4^20^*, *PSBD* and *PSBA* by *ATSIG5* ^18,21^.

Among the six sigma factors in *Arabidopsis*, only *ATSIG5* expression is stress induced and phylogenetically specific^22−24^. It could be induced by high light, low temperature, high salt and osmotic^18^, as well as blue light^22^. Beside these stresses, *MpSIG5* of liverwort *Marchantia polymorpha* is significantly induced by ROS stress^25^. ATSIG5 regulates the renewing capacity from the injury to the photosystem (PS) II reaction center via determining the promoter recognition specificity of PEP in plastid gene expression that activate *PSBD* from the blue-light responsive promoter^10,26^. In addition, ATSIG5 regulates chloroplast *PSBD* and *PSBA* for the PS II core proteins D1 and D2 in response to light quality and intensity, and combines extrinsic and intrinsic signals important for adjusting nuclear and plastid gene transcription in light acclimation processes^27^.

Coordinating the transcription between nuclear encoded photosynthetic genes (NEPGs) and plastid encoded genes (PEGs) to maintain the proper stoichiometry of nuclear encoded proteins, plastid proteins, carotenoids and chlorophylls, is critical for assembly of functional photoprotective and photosynthetic complexes in chloroplasts under stress conditions^28^. Although many abiotic stress-responsive transcription factors (TFs) have been studied, very few are known to modulate the expression of photosynthesis-related genes^6^.

The HD-Zip (homeodomain leucine-zipper) TFs are the most abundant group of homeobox genes expressed only in plants, which have diverse functions during plant development and stress adaptation^29-35^. According to their distinctive features such as gene structures, DNA-binding specificities, additional common motifs and physiological functions, HD-Zip TFs can be classified into four subfamilies^36^. There are 10 HD-Zip II genes in *Arabidopsis* genome, which play important roles from auxin response to shade avoidance. Five HD-Zip class II genes, including *HAT1*, *HAT2*, *HAT3*, *ATHB2*, *ATHB4*, are known to respond to light quality changes^37^. Auxin response analyses strongly suggested that *HAT1, HAT3* and *ATHB4* were under the control of the phytochrome system as *ATHB2* ^38^. For the remaining class II HD-Zip members, little is known about their functions except ATHB17.

ATHB17 localizes to both the cytoplasm and nuclei, which is regulated by its unique N-terminus. Overexpression of ATHB17 in *Arabidopsis* enhances chlorophyll content in the leaves, while expression of a truncated ATHB17 protein in maize increases ear weight at silking^39,40^. *ATHB17*-overexpressing *Arabidopsis* plants are sensitive to ABA and NaCl, whereas *ATHB17* knockout mutants are insensitive to ABA and NaCl at post germination stage. However, these phenotypes are weak and the expression of ABA-responsive genes is not significantly altered in the *ATHB17*-overexpressing plants compared with wide-type (WT) plants^41^. Thus, it remains obscure that the phenotypes were resulted from modulating ABA signaling or other mechanisms.

In this study, we find that ATHB17 plays as an important regulator to coordinate expression of NEPGs and PEGs to cope with environmental stresses. *ATHB17* responds to multiple abiotic stresses. Overexpression of *ATHB17* enhances plant tolerance to salt, drought and oxidative stresses, and knockout *ATHB17* results in the opposite phenotypes. By RNA-seq profile analysis, we find ATHB17 represses the expression of many NEPGs while activates the transcription of many PEGs. Further analysis reveals that ATHB17 can directly bind to the promoter of several NEPGs and likely regulate their expression. Meanwhile, ATHB17 can directly bind to *ATSIG5* promoter to activate *ATSIG5* expression to regulate PEGs expression. Our study reveals a novel pathway involving photosynthesis that requires the coordination between NEPGs and PEGs expression in response to multiple abiotic stresses.

## Results

### *ATHB17* is preferentially expressed in roots and responsive to multiple stress signals

To reveal the expression pattern of *ATHB17*, we analyzed transgenic plants
harboring the *ATHB17* promoter-GUS reporter construct (*pATHB17::GUS*). Strong GUS activity was detected in roots of seedlings at different ages (Fig. 1A-a to A-e and A-g). *ATHB17* was also expressed in rosette leaves with much higher expression levels in the leaf veins (Fig. 1A-f). However, at mature stage, *ATHB17* is mainly expressed in root (Fig. 1A-g). There is only weak expression in other organs, such as rosette leaf (Fig. 1A-h), cauline leaf (Fig. 1A-i), flower and young silique (Fig. 1A-j), and mature silique (Fig. 1A-l). These results were further confirmed by quantitative real-time reverse transcription PCR (qRT-PCR) analysis shown in Fig. 1B.

**Figure 1.**
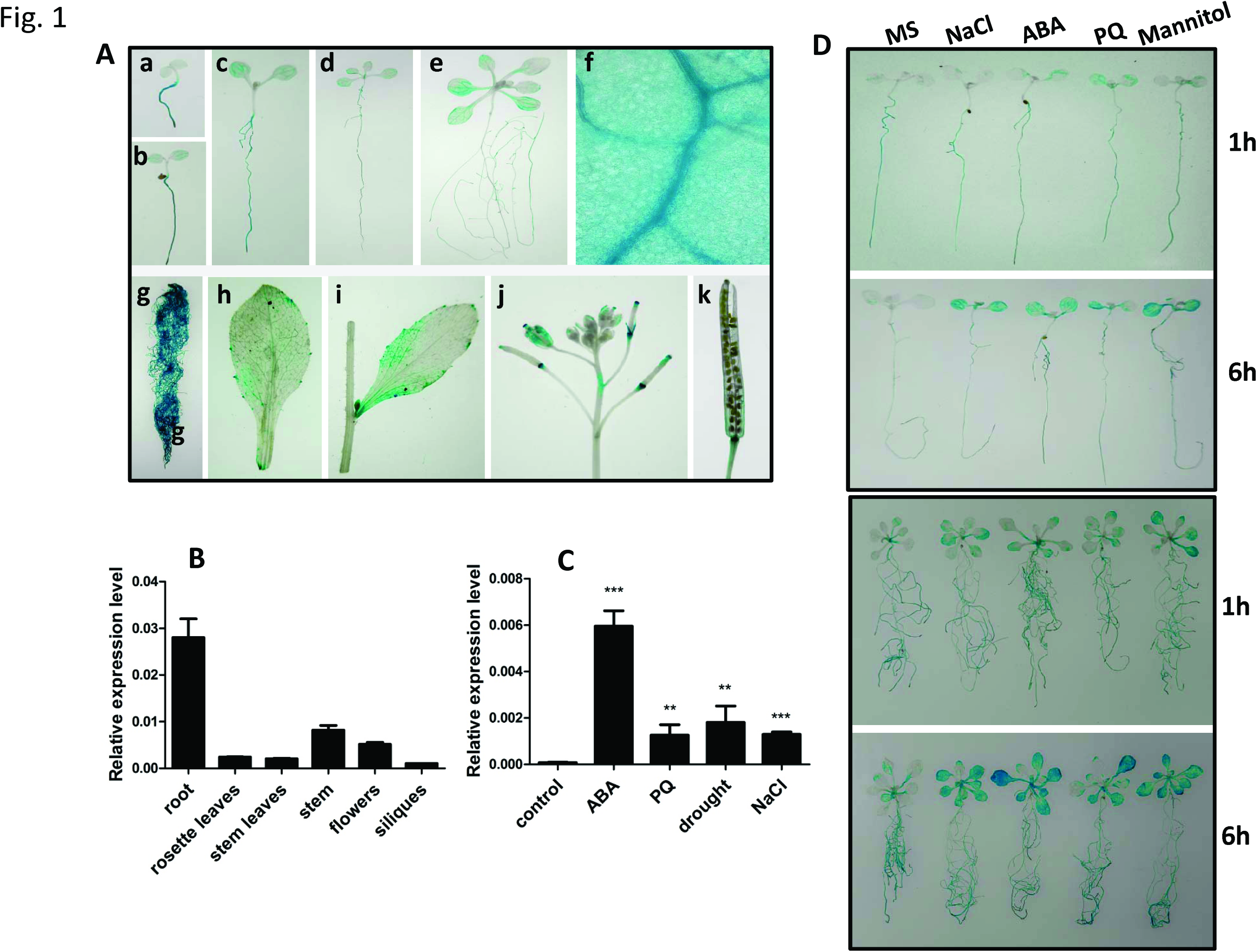
Expression pattern of *ATHB17* and response to different stress signals. (A) Expression pattern of *ATHB17* revealed by GUS staining of *pATHB17::GUS* transgenic plants. GUS activity was observed in seedling of 2-day-old (a), seedling of 4-day-old (b), seedling of 7-day-old (c), seedling of 9-day-old (d), seedling of 14-day-old (e), leaf of 9-day-old plant (f), root of mature plant (g), rosette leaf (h), cauline leaf (i), flower and young silique (j), old silique (k). (B) Analysis of the ATHB17 expression pattern by qRT-PCR. UBQ5 was used as an internal control. Values are mean ± SD of three replica experiments. (C) Expression levels of *ATHB17* after different treatments. Induction levels of *ATHB17* in 10-day-old plants by ABA (100 µM, 4h), PQ (5 µM, 4h), drought (4h), NaCl (200 mM, 4 h) were determined by qRT-PCR. Values are mean ± SD of three replica experiments. (D) Effects of different stress treatments on *pATHB17::GUS* expression. 7-day-old or 14-day-old *pATHB17::GUS* transgenic seedlings grown on MS medium were transferred to MS liquid medium or MS liquid medium containing NaCl (200 mM), ABA (10 µM), PQ (5 µM), mannitol (200 mM) for 1-6 h and then the seedlings were harvested for GUS staining for 3 h.

Moreover, we found that ATHB17 was significantly induced by ABA (abscisic acid), paraquat (PQ), drought, and NaCl treatments (Fig. 1C). Consistent with this result, GUS staining of the *pATHB17::GUS* reporter line treated with NaCl, ABA, PQ, and mannitol also showed strongly induced expression of *ATHB17*, especially in the leaves (Fig. 1D).

### ATHB17 is a positive regulator of tolerance to multiple abiotic stresses

In order to uncover the functions of ATHB17, we generated *35S:: ATHB17* (OX) transgenic *Arabidopsis* plants, and obtained an *ATHB17* knockout (KO) mutant from ABRC (Supplementary Fig. S1). To test whether *ATHB17* is involved in salt tolerance, we determined the NaCl sensitivity of the *ATHB17* OX and KO lines. Firstly, the NaCl sensitivity at germination and seedling establishment stage was assayed. Seeds were germinated and grown in MS (Murashige and Skoog) medium containing different concentration of NaCl. The results show that the *ATHB17* OX lines had a better root growth advantage compared with the WT plants under the stress condition. In contrast, the KO plants showed significantly reduced root growth. However, there was no difference in germination or root elongation on MS medium without NaCl (Fig. 2A). The results of root growth assay indicate that the *ATHB17* KO was more sensitive to NaCl, while the *ATHB17* OX was more tolerant to NaCl than WT at germination and seedling establishment stage (Fig. 2B).

**Figure 2.**
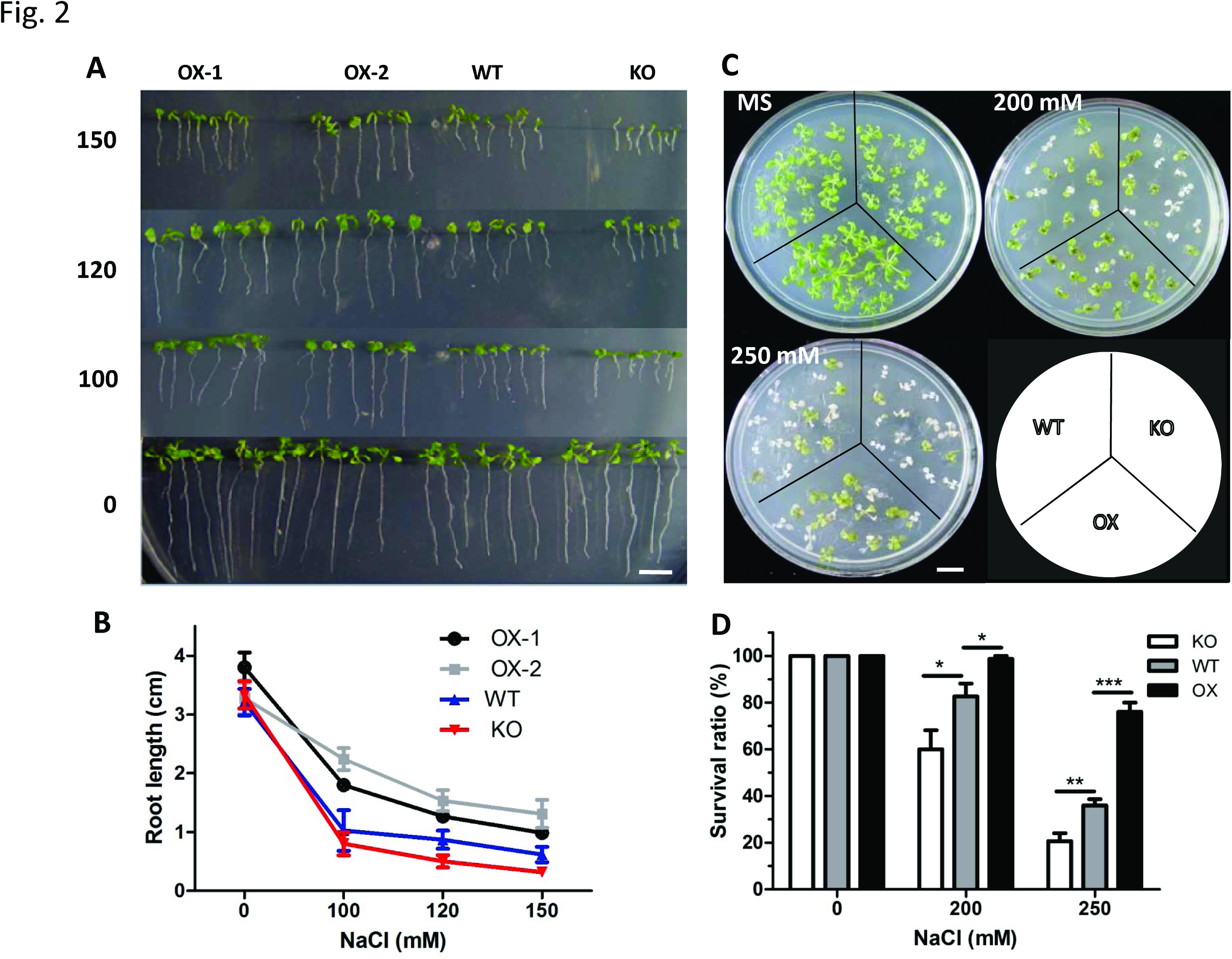
ATHB17 is a positive regulator for salt stress resistance. (A-B) Salt tolerance assay of *ATHB17* OX and KO. Phenotypes of seedlings (A) and root length (B) of *ATHB17* OX, KO and WT plants grew on medium containing different concentrations of NaCl. Seeds were germinated and grew vertically on MS medium or MS medium containing different concentrations of NaCl for 10 days. Bar = 1 cm. (C) Phenotype of *ATHB17* OX, KO and WT plants on MS medium with different concentrations of NaCl. 7-day-old plants on MS medium were transferred to MS medium or MS medium containing different concentrations of NaCl for 7 days. Bar = 4 cm. (D) Survival ratio of the *ATHB17* OX, KO and WT plants on MS medium or MS medium with different concentrations of NaCl.

In another salt tolerance assay, 7-day-old *ATHB17* OX, KO and WT plants were transferred to MS medium or MS medium containing NaCl. After another 7 days growth, the *ATHB17* OX plants showed higher survival ratio whereas the *ATHB17* KO plants showed opposite phenotypes compared with the WT (Fig. 2C-D).

To confirm this, we conducted complementation analysis by expressing *ATHB17* in the *ATHB17* KO plants, which restored the salt tolerance of KO plants to wild-type level (Supplementary Fig. S2). Moreover, *ATHB17* was able to confer salt tolerance in tobacco when overexpressed (Supplementary Fig. S3).

Since *ATHB17* is responsive to multiple stresses as demonstrated above, we also analyzed the role of *ATHB17* in drought stress and oxidative stress. The *ATHB17* OX *Arabidopsis* plants showed enhanced drought tolerance while the *ATHB17* KO plants showed drought sensitive compared with WT plants (Supplementary Fig. S4A-B), which is consistent with the result of Park et al.^41^. In addition, overexpression of *ATHB17* also conferred improved tolerance to PQ (Supplementary Fig. S4C-D). These data indicate ATHB17 is a positive regulator of multiple abiotic stresses.

### The expression of typically stress-responsive marker genes was not significantly affected by ATHB17

To investigate the mechanism of ATHB17 modulating stress tolerance, we compared the transcriptomes of *ATHB17* OX, KO and WT plants grown under normal and 200 mM NaCl treated conditions using RNA-seq method. Surprisingly, we found the expression of classic stress-induced marker genes such as *RD29A*, *RD29B*, *RD20*, *RD22*, *COR47*, *COR45B*, *CBF1*, *SOS2*, *SOS3*, were not significantly influenced by *ATHB17* both under normal and salt treated conditions. Expression levels of ABA signaling and synthesis pathway genes were also not significantly changed in the *ATHB17* OX and KO plants (Table 1). To validate the results of RNA-seq profiling analysis, eight stress-responsive genes were selected for qRT-PCR analysis. The results showed that the response of these genes was not significantly affected by the expression level of *ATHB17*. Taken together, these results implicate that *ATHB17*-conferred stress tolerance is probably not through the classical stress response pathway but through a new pathway.

**Table 1.** Relative expression levels of stress-responsive genes identified by RNA-seq profiling.

| Gene ID | Annotation | Normal (TPM) |  |  | NaCl (TPM) |  |  |
| --- | --- | --- | --- | --- | --- | --- | --- |
|  |  | WT | OX | KO | WT | OX | KO |
| AT5G52310 | RD29A (responsive to desiccation 29A) | 0.99 | 0.38 | 0.01 | 71.54 | 73.85 | 52.09 |
| AT5G52300 | RD29B (responsive to desiccation 29B) | 0.01 | 0.01 | 0.28 | 16.32 | 14.61 | 12.21 |
| AT5G25610 | RD22 (responsive to desiccation 22) | 36.67 | 11.53 | 44.81 | 182.8 | 280.38 | 373.55 |
| AT2G33380 | RD20 (responsive to desiccation 20) | 16.52 | 20.36 | 10.65 | 116.37 | 146.30 | 133.47 |
| AT1G20440 | COR47 (cold regulated 47) | 943.44 | 770.26 | 1405.98 | 3953.93 | 4602.60 | 5296.48 |
| AT2G42530 | COR15B (cold regulated 15B) | 0.01 | 0.01 | 0.01 | 4.12 | 3.80 | 6.10 |
| AT4G25490 | CBF1 (C-repeat/DRE binding factor 1) | 0.01 | 0.01 | 0.01 | 15.49 | 20.61 | 19.94 |
| AT4G26080 | ABI1 (abscisic acid insensitive 1) | 36.01 | 46.48 | 44.95 | 122.14 | 122.88 | 186.78 |
| AT5G57050 | ABI2 (abscisic acid insensitive 2) | 0.66 | 5.57 | 2.63 | 74.01 | 64.04 | 61.45 |
| AT2G36270 | ABI5 (abscisic acid insensitive 3) | 3.96 | 3.07 | 4.29 | 17.47 | 23.62 | 16.68 |
| AT5G67030 | ABA1 (abscisic acid deficient 1) | 49.55 | 44.95 | 75.24 | 287.96 | 282.38 | 264.50 |
| AT1G16540 | ABA3 (abscisic acid deficient 3) | 8.26 | 17.86 | 8.71 | 24.23 | 35.42 | 17.09 |
| AT3G14440 | NCED3 (9-cis-epoxycarotenoid dioxygenase 3) | 0.99 | 0.38 | 0.01 | 71.54 | 73.85 | 52.09 |
| AT5G35410 | SOS2 (salt overly sensitive 2) | 9.58 | 10.95 | 6.64 | 7.75 | 10.21 | 7.32 |
| AT5G24270 | SOS3 ( salt overly sensitive 3) | 1.32 | 4.03 | 1.38 | 0.82 | 0.40 | 0.01 |

TPM: number of transcripts per million tags.

### ATHB17 negatively regulates some NEPGs while positively regulates many PEGs

Based on Gene Ontology (GO) term enrichment analysis of the RNA-seq profiling data, we found the NEPGs were significantly enriched among the different expressed genes between the ATHB17 OX and ATHB17 KO both under normal and salt stress conditions (Supplementary Data 1 and 2). The expression of 26 of the 63 NEPGs in the *Arabidopsis* genome were found down-regulated (> 2 fold) in the *ATHB17* OX compared with *ATHB17* KO under normal or NaCl treated conditions. More specifically, under normal condition, 20 of the 26 NEPGs had reduced expression (> 1.5 fold) in the *ATHB17* OX plants compared with *ATHB17* KO, with 16 NEPGs up-regulated in the *ATHB17* KO while down-regulated in the *ATHB17* OX compared with WT.

Under salt treated condition, 21 of the 26 NEPGs were detected down-regulated expressing in the *ATHB17* OX compared with *ATHB17* KO plants, with 16 NEPGs had higher expression in *ATHB17* KO while lower expression in *ATHB17* OX (Table 2). These genes have functions in PSI complexes (*PSAF*, *PSAH2*, *PSAH1*, *PSAD1*, *PSAG*), light-harvesting complexes (*LHCA1, LHCA2*, *LHCA3, LHCA4*, *LHCA5, LHCB3*, *LHCB5*, *LHCB6*, *LHCB7, LHB1B1*, *LHB1B2*), chlorophyll a/b-binding (*CAB1*, *CAB3*) proteins, and PS II oxygen evolving complex (*PSBO1*, *PSBO2*, *PPL1*, *PPL2*).

**Table 2.** Expression levels of NEPGs in *ATHB17* OX, KO and WT plants under normal or NaCl treated conditions identified by RNA-seq profiling

| Gene ID | Description | Normal (TPM) |  |  | NaCl (TPM) |  |  |
| --- | --- | --- | --- | --- | --- | --- | --- |
|  |  | KO | WT | OX | KO | WT | OX |
| AT1g29930 | CAB1 (chlorophyll a/b binding protein 1) | 2659.41 | 1921.89 | 172.11 | 5531.28 | 4153.04 | 1530.00 |
| AT1g31330 | PSAF (PS I subunit F) | 1117.89 | 912.05 | 284.09 | 1296.45 | 946.94 | 291.59 |
| AT2g34420 | LHB1B2 (light-harvesting chlorophyll protein complex II subunit B2) | 2781.39 | 2899.35 | 712.06 | 4826.90 | 4362.05 | 1421.93 |
| AT3g55330 | PPL1 (PsbP-like protein 1) | 37.48 | 14.53 | 5.57 | 29.30 | 17.14 | 9.41 |
| AT3g61470 | LHCA2 (PS I light harvesting complex gene 2) | 2269.95 | 2062.94 | 705.34 | 4063.92 | 3235.44 | 1422.93 |
| AT4g10340 | LHCB5 (light harvesting complex of PS II subunit 5) | 836.73 | 592.29 | 86.05 | 1080.78 | 707.61 | 357.03 |
| AT4g30950 | FAD6 (fatty acid desaturase 6) | 182.28 | 154.60 | 67.61 | 260.43 | 212.79 | 127.28 |
| AT5g66570 | PSBO1 (PS II oxygen-evolving complex 1) | 2749.86 | 2251.56 | 485.78 | 4219.77 | 3937.28 | 1953.87 |
| AT1g52230 | PSAH2 (PS I subunit H2) | 69.84 | 92.16 | 26.70 | 72.84 | 60.00 | 30.22 |
| AT2g34430 | LHB1B1 (light-harvesting chlorophyll protein complex II subunit B1) | 1449.82 | 1655.64 | 306.57 | 4226.69 | 3826.35 | 2079.15 |
| AT3g16140 | PSAH1 (PS I subunit H1) | 143.97 | 158.89 | 57.24 | 196.95 | 166.81 | 48.03 |
| AT1g15820 | LHCB6 (light harvesting complex of PS II subunit 6) | 1560.05 | 1324.97 | 614.10 | 1194.31 | 1474.56 | 735.88 |
| AT1g29910 | CAB3 (chlorophyll a/b binding protein 3) | 821.65 | 702.62 | 110.45 | 1632.16 | 1800.76 | 510.13 |
| AT2g39470 | PPL2 (PsbP-like protein 2) | 46.05 | 21.47 | 15.17 | 63.48 | 79.12 | 26.82 |
| AT3g47470 | LHCA4 (light-harvesting chlorophyll protein complex I subunit A4) | 1025.23 | 863.49 | 428.16 | 1665.12 | 2171.46 | 1119.13 |
| AT3g50820 | PSBO2 (PS II subunit O-2) | 146.05 | 87.21 | 46.29 | 316.18 | 454.27 | 462.50 |
| AT3g54110 | PUMP1 (plant uncoupling mitochondrial protein 1) | 87.13 | 71.68 | 40.53 | 43.95 | 49.28 | 64.84 |
| AT4g02770 | PSAD1 (PS I subunit D-1) | 2057.38 | 1748.79 | 900.69 | 1283.02 | 1341.55 | 856.36 |
| AT5g54270 | LHCB3 (light-harvesting chlorophyll B-binding protein 3) | 294.44 | 231.23 | 70.69 | 320.65 | 348.94 | 127.28 |
| AT1g61520 | LHCA3 (PS I light harvesting complex gene 3) | 2178.95 | 1902.07 | 1325.00 | 3721.29 | 2438.82 | 1668.09 |
| AT1g45474 | LHCA5 (PS I light harvesting complex gene 5) | 55.87 | 67.72 | 31.50 | 46.39 | 35.77 | 20.21 |
| AT1g55670 | PSAG (PS I subunit G) | 37.48 | 37.33 | 23.63 | 38.66 | 24.56 | 13.81 |
| AT1g67740 | PSBY (PS II BY) | 89.90 | 63.75 | 142.91 | 135.10 | 154.12 | 64.44 |
| AT1g76570 | LHCB7 (light-harvesting complex B7) | 8.44 | 7.93 | 9.03 | 24.82 | 11.21 | 10.81 |
| AT3g54890 | LHCA1 (PS I light harvesting complex gene 1) | 53.38 | 122.55 | 222.82 | 103.76 | 94.78 | 50.03 |
| AT4g28660 | PSB28 (PS II reaction centre PSB28 protein) | 33.33 | 10.90 | 20.75 | 13.43 | 10.71 | 4.00 |

qRT-PCR analysis was carried out to validate the results of expression profiling analysis. The data in Fig 4A showed that 11 of the 13 tested NEPGs had lower transcript levels in *ATHB17* KO plants while had higher expression levels in *ATHB17* OX plants compared with WT plants. This result is consistent with the data of RNA-seq profiling analysis. These data indicate that ATHB17 may plays as a repressor of NEPGs, which agrees with its function as a repressor reported by Rice et al.^40^.

**Figure 3.**
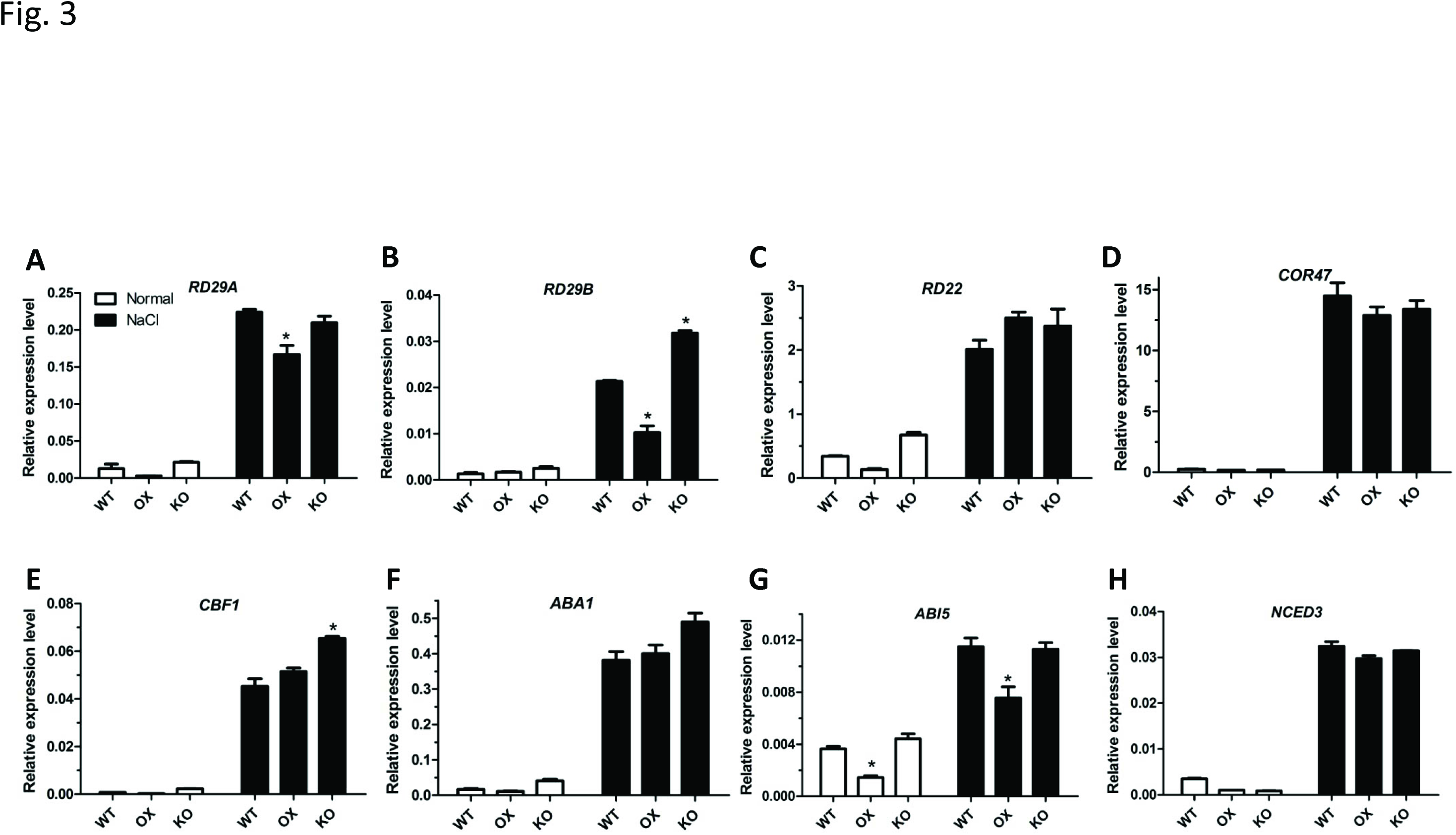
qRT-PCR validation of the data of stress-responsive marker genes in the RNA-seq profiling. 12-day old seedlings of the *ATHB17* OX, KO lines and WT plants were treated with liquid MS medium containing 0 or 200 mM NaCl for 5 h. Total RNA were extracted and reverse-transcribed as templates for qRT-PCR. UBQ5 was used as an internal control. Values are mean ± SD of three replica experiments (*P < 0.05, ***P < 0.001).

**Figure 4.**
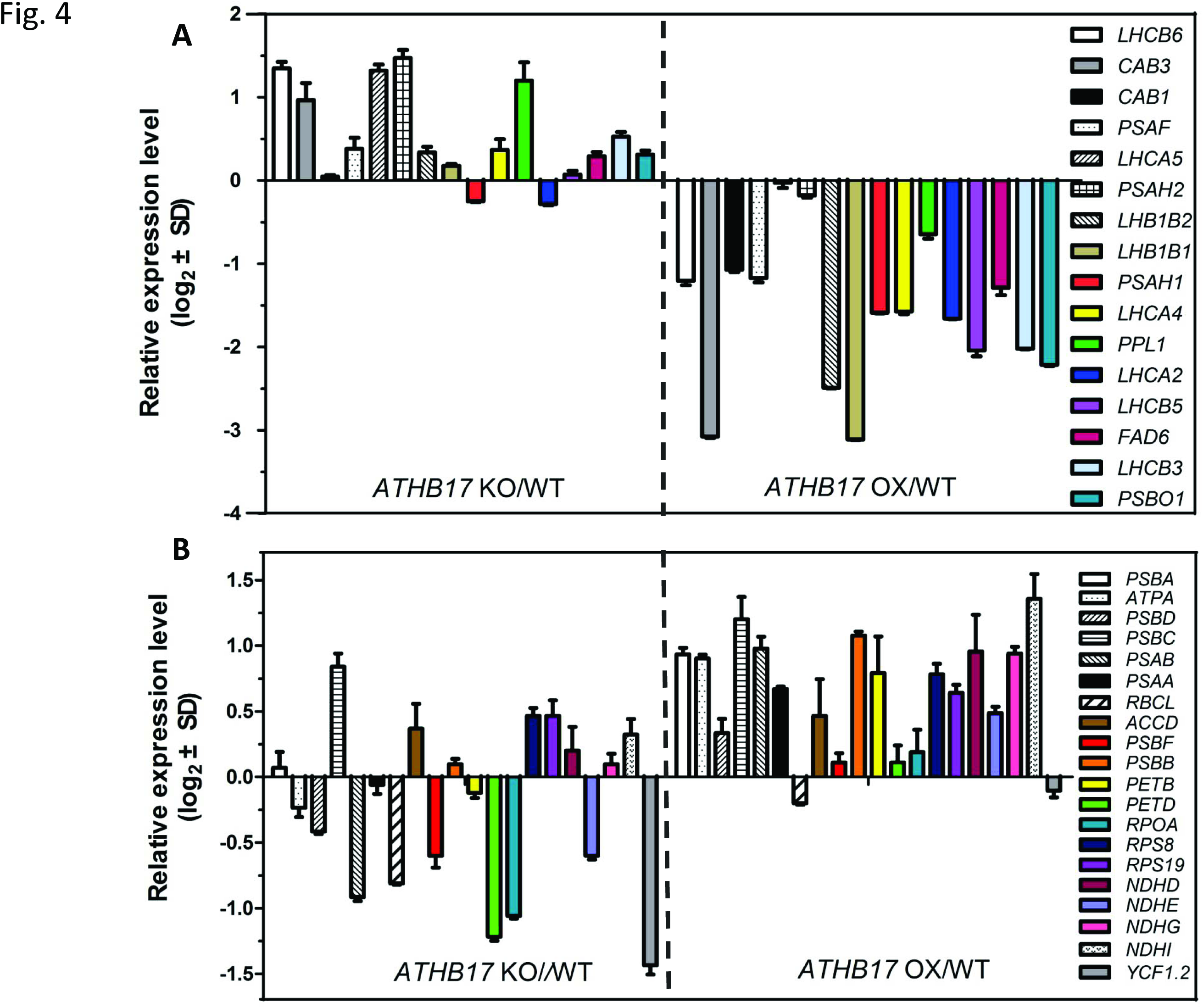
qRT-PCR validation of the expression level of genes encoded by both the nucleus genome and chloroplast genome in the profiles of RNA-seq analysis results. 12-day-old plants grown under normal condition were used for qRT-PCR analysis. Relative transcription levels of the genes in *ATHB17* OX and KO are normalized to levels in WT control (WT = 0). (A) The expression levels of NEPGs are down-regulated by *ATHB17.* Values are mean ± SD of three independent experiments. (B) The expression levels of PEGs are positively affected by *ATHB17.* Values are mean ± SD of three independent experiments.

Interestingly, we found that the expression of many PEGs is positively correlated with *ATHB17* expression level. 20 of the 35 detected PEGs had markedly higher expression in the *ATHB17* OX plants and lower expression in the *ATHB17* KO plants compared with WT plants under normal or salt treated conditions (Table 3). These genes mainly belong to PSIcomplexes *(PSAA, PSAB)*, PSIIcomplexes (*PSBD*, *PSBC*, *PSBB*, *PSBA*, *PSBF*), NAD(P)H dehydrogenase *(NDHD, NDHI, NDHG, NDHE)*, cytochrome b6f complexes *(PETB, PETD).* As shown in Table 3, transcript of 15 of the 20 PEGs had increased in *ATHB17 OX* plants while reduced in *ATHB17* KO plants compared with WT plants under normal condition. Moreover, 13 of the 20 PEGs in *ATHB17 OX* plants were up-regulated compared with *ATHB17 KO* and WT plants after salt treatment. However, no significant expression changes of these genes between *ATHB17 KO* and WT plants were found under salt treated condition.

**Table 3.** Expression levels of PEGs in *ATHB17* OX, KO and WT plants under normal or NaCl treated conditions identified by RNA-seq profiling

| Gene ID | Description | Normal (TPM) |  |  | NaCl (TPM) |  |  |
| --- | --- | --- | --- | --- | --- | --- | --- |
|  |  | KO | WT | OX | KO | WT | OX |
| ATCG00270 | PSBD (PS II D2 protein) | 3.73 | 5.62 | 9.99 | 8.55 | 7.91 | 20.61 |
| ATCG00280 | PSBC (chloroplast gene encoding a CP43 subunit of the PSII reaction center) | 18.53 | 31.05 | 36.50 | 46.39 | 42.53 | 99.06 |
| ATCG00340 | PSAB (encodes the D1 subunit of PS I and II reaction centers) | 45.22 | 98.77 | 106.42 | 89.93 | 80.27 | 215.54 |
| ATCG00350 | PSAA (encodes psaa protein comprising the reaction center for PS I) | 21.44 | 62.10 | 70.88 | 47.61 | 49.28 | 103.07 |
| ATCG00680 | PSBB (encodes for CP47, subunit of the PS II reaction center) | 13.14 | 16.52 | 19.78 | 17.09 | 13.19 | 32.02 |
| ATCG00740 | RPOA (RNA polymerase alpha subunit) | 8.16 | 21.14 | 22.09 | 8.55 | 14.50 | 49.03 |
| ATCG00820 | RPS19 (encodes a 6.8-kDa protein of the small ribosomal subunit) | 1.38 | 1.98 | 2.69 | 1.22 | 0.82 | 5.20 |
| ATCG01050 | NDHD (represents a plastid-encoded subunit of a NAD(P)H dehydrogenase complex) | 1.11 | 4.62 | 9.80 | 1.22 | 0.99 | 3.20 |
| ATCG01090 | NDHI (encodes subunit of the chloroplast NAD(P)H dehydrogenase complex) | 0 | 0.66 | 1.73 | 0 | 0.99 | 1.20 |
| ATCG00730 | PETD (a chloroplast gene encoding subunit IV of the cytochrome b6/f complex) | 0.97 | 0.99 | 1.34 | 1.22 | 0.33 | 3.20 |
| ATCG01080 | NDHG (NADH dehydrogenase ND6 ) | 0.55 | 1.65 | 3.07 | 1.63 | 0.33 | 2.20 |
| ATCG00020 | PSBA (encodes chlorophyll binding protein D1, a part of the PS II reaction center core) | 18.12 | 31.71 | 68.96 | 36.22 | 27.69 | 36.02 |
| ATCG00490 | RBCL (large subunit of RUBISCO) | 73.44 | 106.7 | 110.64 | 141.20 | 87.52 | 139.29 |
| ATCG00570 | PSBF (PS II cytochrome b559) | 0 | 1.98 | 2.11 | 1.22 | 2.47 | 1.60 |
| ATCG00720 | PETB (encodes the cytochrome b6 subunit of the cytochrome b6/f complex) | 0.97 | 1.98 | 4.80 | 3.26 | 1.65 | 2.80 |
| ATCG01130 | YCF1.2 (hypothetical protein) | 1.94 | 4.62 | 9.22 | 2.44 | 5.44 | 4.20 |
| ATCG01070 | NDHE (NADH dehydrogenase ND4L) | 0 | 0 | 0.96 | 0 | 0.82 | 0 |
| ATCG00500 | ACCD (encodes the carboxytransferase beta subunit of the Acetyl-CoA carboxylase complex in plastids) | 0 | 0.66 | 0.38 | 0 | 0 | 0.60 |
| ATCG00770 | RPS8 (chloroplast 30S ribosomal protein S8) | 0 | 1.65 | 0.77 | 0 | 0.49 | 0.60 |
| ATCG00120 | ATPA (encodes the ATPase alpha subunit) | 0.97 | 2.97 | 2.31 | 1.22 | 1.32 | 1.20 |

RT-PCR validation showed a similar expression pattern of these genes as RNA-seq profiling analysis. Most of the genes tested had higher expression in *ATHB17 OX* and relatively lower expression in *ATHB17 KO* as showed in Fig. 4B. Taken together, our data imply that ATHB17 positively regulates the expression of many PEGs.

### ATHB17 can directly bind to the promoters of several NEPGs

HD-ZIP transcription factors show a binding preference for variant HD-binding sequences^36,42^. According to the result of bioinformatic analysis, combined with the information from AGRIS *(Arabidopsis* Gene Regulatory Information Server), we selected six kinds of 8-9 bp sequences as potential HD binding sites. We firstly tested the binding affinities of ATHB17 to these sequences by yeast one-hybrid (Y1H) assay. We found full length ATHB17 protein had strong self-activation activity. It had reported that ATHB17 protein lacking the first 113 amino acids (ATHB17Δ113) can still homodimerize and specifically recognize the sequence requirements for DNA binding^40^, thus we used ATHB17 protein lacking the first 107 amino acids (ATHB17Δ107) for Y1H and mobility shift assay (EMSA). As results showed in Supplementary Fig. S5A, ATHB17Δ107 had no self-activation activity. Y1H assay reveled ATHB17 had strong binding affinities to these HD binding cis-elements: aaattagt, tttaattt and taaatgta (Supplementary Fig. S5B). Then we did EMSA to confirm the binding affinities of ATHB17 to the potential ATHB17 binding cis-elements *in vitro*. Supplementary Fig. S5C shows that ATHB17 could directly bind to the sequences aaattagt and tttaattt, but failed to bind to taaatgta *in vitro*. The results of EMSA indicate that ATHB17 protein was able to directly bind to the tttaattt motif *in vitro*, and the binding was specific as demonstrated by competition assay using unlabelled (competitor) and non-specific probes (non-competitor) (Supplementary Fig. S5D).

Based on the above analysis, we chose aaattagt and tttaattt as the ATHB17 binding cis-elements. Promoter sequence analysis of the 19 ATHB17 down-regulated NEPGs revealed that 10 of the 19 genes had at least one ATHB17 binding cis-element (Fig. 5A). To analyze whether ATHB17 could directly regulate expression of all these 10 genes by binding to the different ATHB17 cis-elements in their promoters, we performed Y1H assay. As shown in Fig. 5B, ATHB17 could only bind to the promoter of 5 NEPGs (*FDA6*, *LHCA2*, *LHB1B1*, *LHB1B2*, *PSBO1)* with different binding affinities. Subsequently, chromatin immunoprecipitation (ChIP) assays using the transgenic plants expressing 35S-haemagglutinin (HA)-*ATHB17* plants were conducted to validate the bind affinities *in vivo*. ChIP-qPCR showed that ATHB17 could directly bind to ATHB17 binding cis-element motifs in the promoters of the 5 genes screened out by Y1H (Fig. 5C). However, ATHB17 failed to bind the h fragment in *PSBO1* promoter, which also had no binging affinity to ATHB17 in yeast. These results are consistent with the data of Y1H and the expression pattern of these genes in the *ATHB17* OX, KO and WT plants (Table 2). These results indicate that ATHB17 can directly bind to the promoter of a number of NEPGs to regulate their expression.

**Figure 5.**
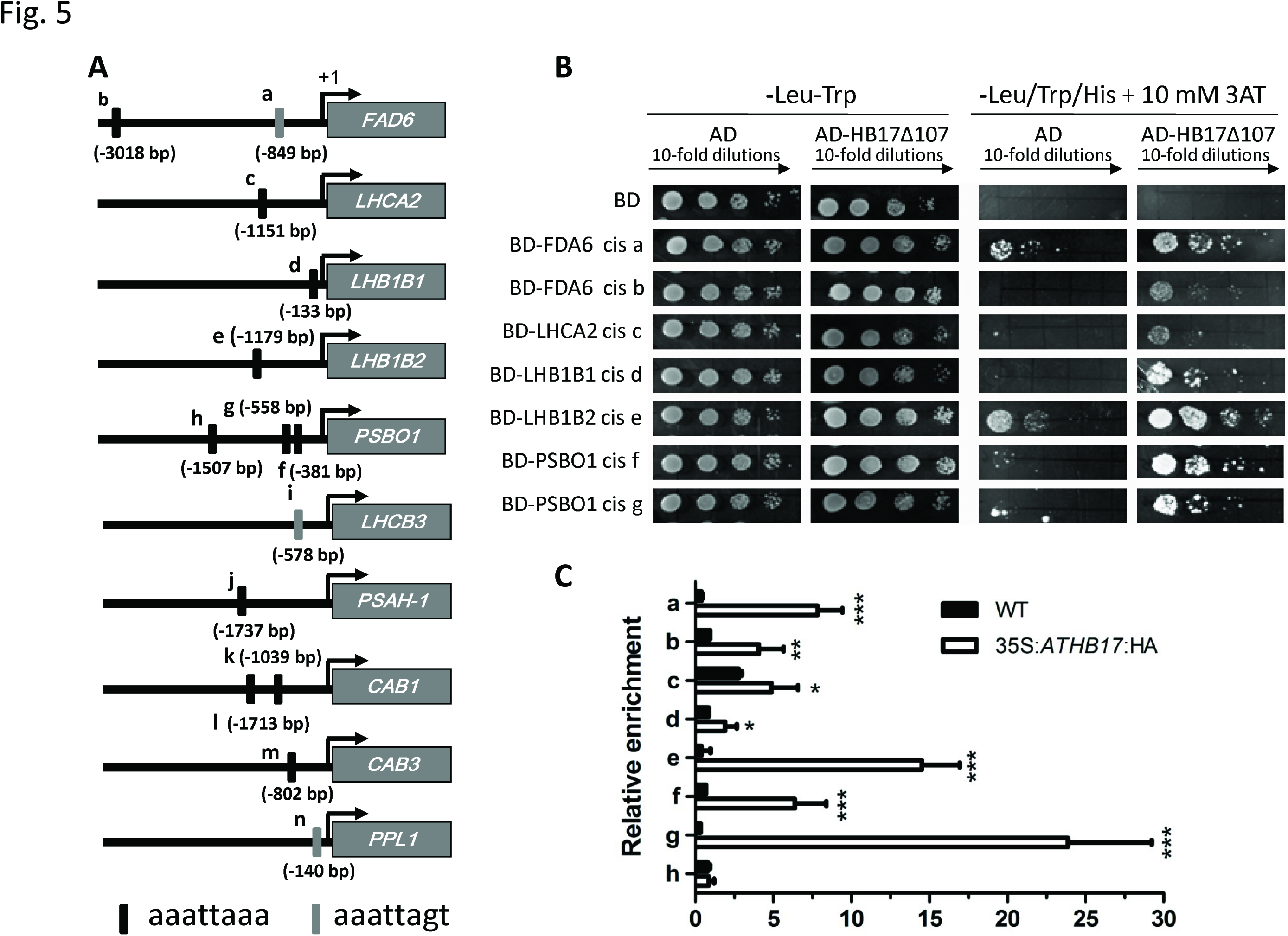
ATHB17 binds to the promoters of several NEPGs. (A) Location of ATHB17 binding cis-elements in the promoter of several NEPGs. The ATHB17 binding cis-elements are indicated with filled rectangles, above/below which the sites of the last base of the cis-elements relative to the start code are shown. (B) Y1H assay for ATHB17 binding to the 25 bp fragment containing ATHB17 binding cis-element from the promoter of five NEPGs, respectively. (C) ChIP assay. About 70-200 bp promoter fragments containing ATHB17 binding cis-element were enriched by anti-HA antibodies in ChIP-qPCR analysis. Values are mean ± SD of three independent experiments (*P < 0.05, **P < 0.01, ***P < 0.001).

### ATHB17 binds to the HD cis-elements in the *ATSIG5* promoter

Through RNA-seq profiling analysis, we found a nuclear encoded sigma factor, *ATSIG5*, had significantly higher expression in *ATHB17* OX plants than in WT and *ATHB17* KO plants both under normal and salt stress conditions (Fig. 6A). qRT-PCR validation showed that *ATHB17* OX plants had increased *ATSIG5* transcription compared with WT and *ATHB17* KO plants under normal condition. After salt treatment, *ATSIG5* transcript was significantly up-regulated in *ATHB17 OX* plants, while down-regulated in *ATHB17 KO* plants compared with WT plants (Fig. 6B). Promoter sequence analysis revealed two potential ATHB17 binding cis-elements in the *ATSIG5* promoter: cis1 tttaattt located 1148 bp upstream of start codon and cis2 aaattagt located 1054 bp upstream of start codon (Fig.6 C). These data suggest that *ATSIG5 may* be a candidate target of ATHB17.

**Figure 6.**
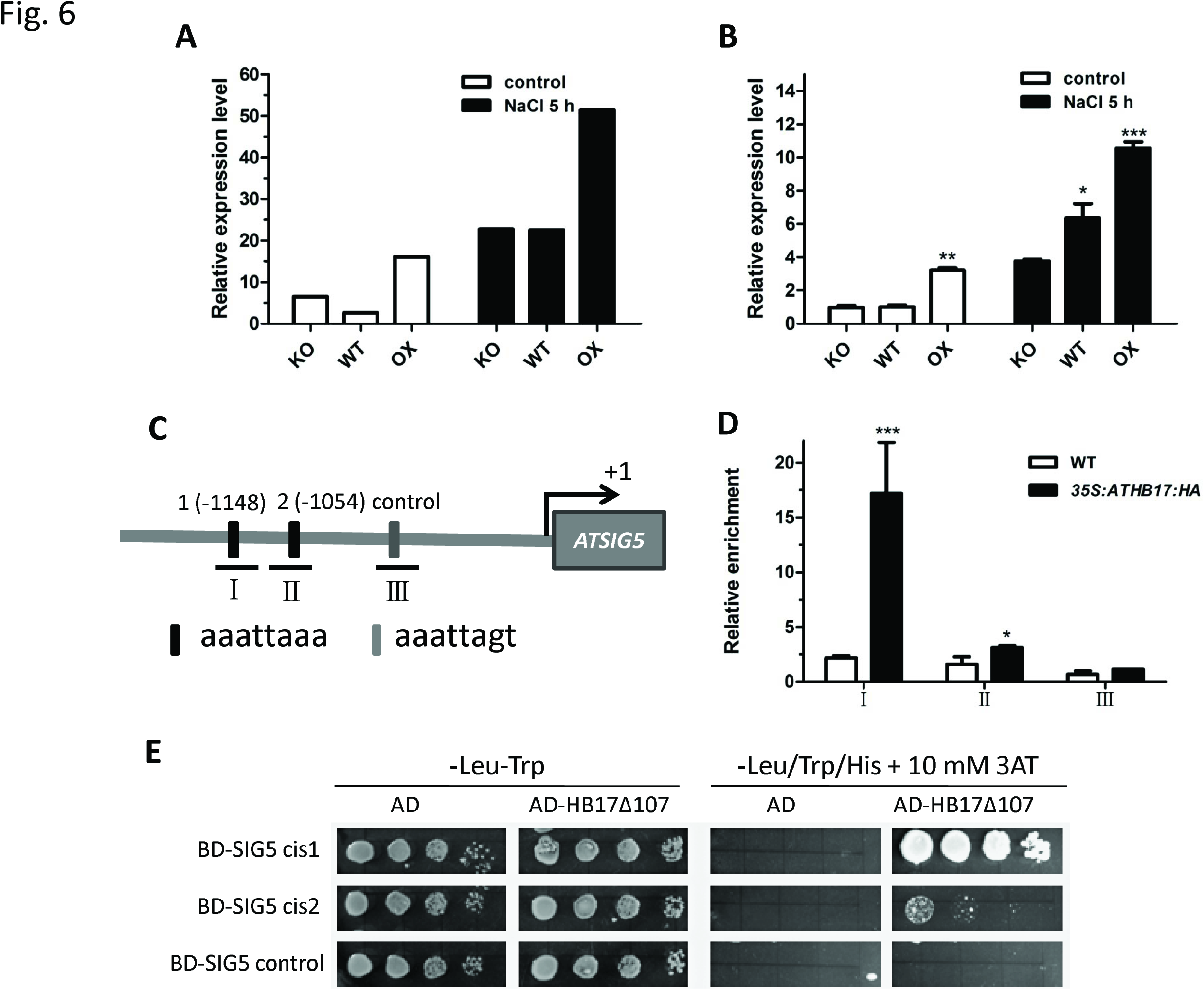
ATHB17 directly regulates the transcription of *ATSIG5*. (A) Expression levels of *ATSIG5* in the RNA-seq profiling data. (B) qRT-PCR validation of the expression levels of *ATSIG5* in RNA-seq profiling. 12-day-old plants grown on MS medium were used transferred to liquid MS medium containing 0 or 200 mM NaCl for 5 h. The plants were harvested for qRT-PCR analysis. Values are mean ± SD and asterisks denote Student’s *t-test* significance compared with KO (\**P* < 0.05, ** *P <* 0.01, *** *P <* 0.001). (C) The schematic illustration of the locations of ATHB17 binding cis-elemnt in the promoters of *ATSIG5 and* the fragments (short lines) used in ChIP-qPCR assay. cis1 is tttaattt located 1148 bp upstream of start codon, cis2 is aaattagt located 1054 bp upstream of start codon, control is a random 8 bp sequence. (D) qPCR data from ChIP assay with antibody against HA. A fragment without ATHB17 binding cis-element (fragment _) was used as negative control. Values are mean ± SD of three replica experiments (*P < 0.05,***P < 0.001). (E) Y1H assay for ATHB17 binding to the 25 bp fragment containing ATHB17 binding cis-element from the promoter of *ATSIG5.* Fragment containing no ATHB17 binding cis-element was used as a negative control.

ChIP and Y1H assay was conducted to determine whether ATHB17 could directly bind to the HD cis-elements in the promoter of *ATSIG5.* The results of ChIP-qPCR showed in Fig. 6D revealed that ATHB17 was able to bind to the *ATSIG5* promoter DNA fragmentland II which containing the HD cis-elements. However, the DNA fragment III containing no HD cis-elements was not enriched, suggesting that ATHB17 specifically binds to the HD cis-element motifs in the *ATSIG5* promoter. Binding of ATHB17 to the *ATSIG5* promoter was further tested by Y1H assay. Consistent with the ChIP assay results, ATHB17 could directly bind to both of the promoter fragments containing HD cis-element cis1 and cis2 in yeast cells, with much stronger binding affinities to the region containing cis1, but could not bind to the promoter fragment without HD cis-elements as a negative control (Fig. 6E). These data indicate that *ATSIG5* was a direct target of ATHB17.

### ATHB17 regulates salt stress tolerance partly by modulating *ATSIG5*

*ATSIG5 was* reported to respond to multiple stress signals, including salt stress. The *sig5-1* mutant is hypersensitive to NaCl treatment^18^. To study the functions of ATSIG5, we generated *35S::ATSIG5* (OX) lines and obtained *ATSIG5* knockout mutants *sig5-1* and *sig5-4* (Supplementary Fig. S6). After germinated and grew vertically on MS or MS medium containing NaCl for 10 days, the *ATSIG5 OX* showed salt tolerant while *sig5-1* and *sig5-4* showed salt sensitive phenotypes compared with the WT (Fig. 7). These results agree with that reported by Nagashima et al.^18^.

**Figure 7.**
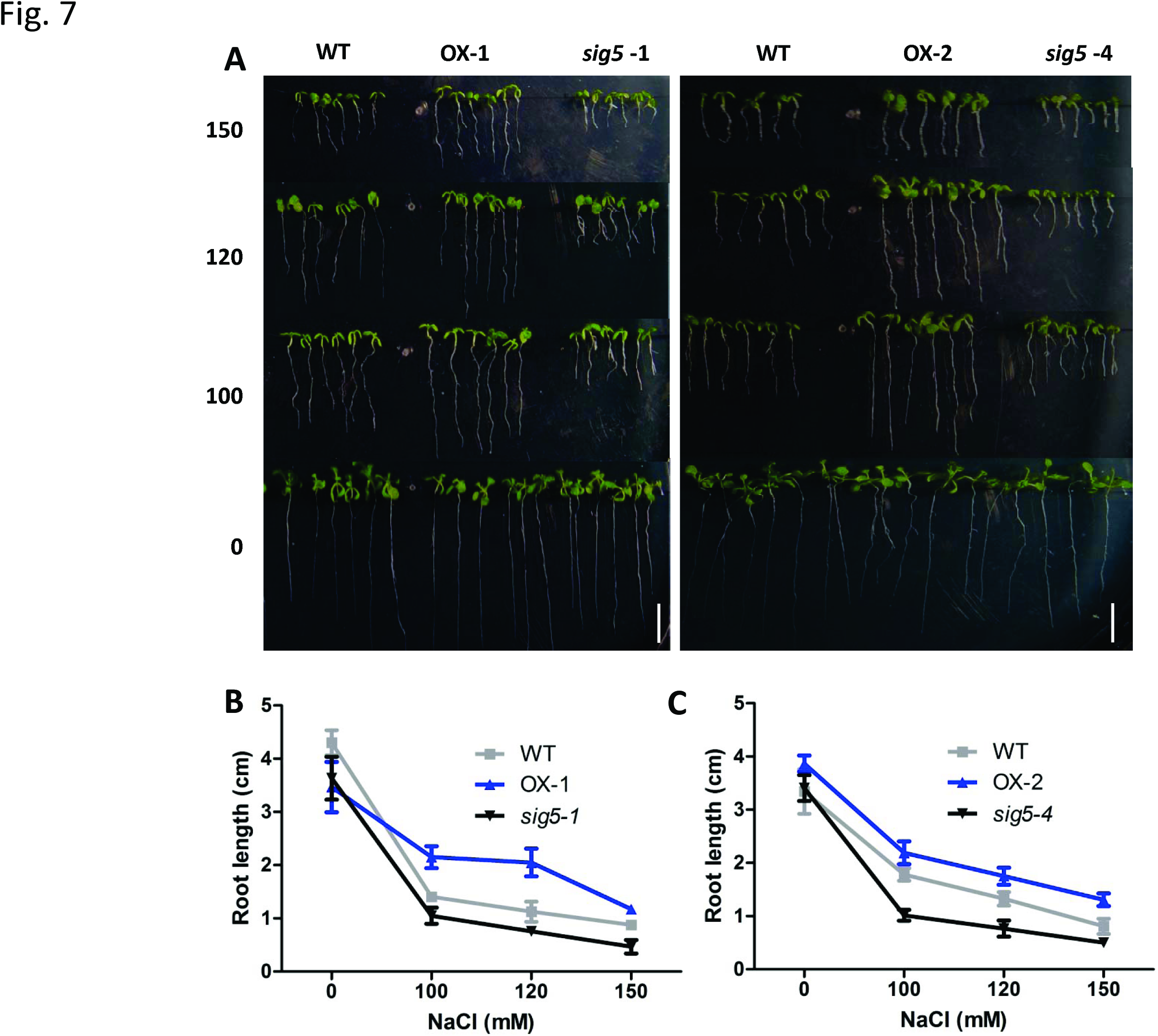
*ATSIG5*-overexpressing transgenic plants were more tolerant while its knockout mutants were more sensitive to salt stress. (A) Salt tolerance assay of *ATSIG5*-overexpressing and knockout plants. Seeds were sowed on MS medium containing 0, 100, 120, 150 mM NaCl and grew vertically for 10 days. Bar = 1 cm. (B-C) Root length of the 10-day-old plants grown on MS medium containing different concentrations of NaCl.

To investigate further whether *ATHB17* modulating salt stress through regulating *ATSIG5* transcript, we introduced *ATHB17* OX into *sig5-1* background by crossing. The *ATHB17* OX *sig5-1* offsprings were tested for salt tolerance. Although *ATHB17* gene was overexpressed in the hybrid offspring, the *ATHB17* OX *sig5-1* seedlings did not show salt stress resistant phenotypes as *ATHB17* OX plants did but an intermediate phenotype between the two parents (Fig. 8), implying that ATHB17-conferred stress tolerance is partially dependent on *ATSIG5*.

**Figure 8.**
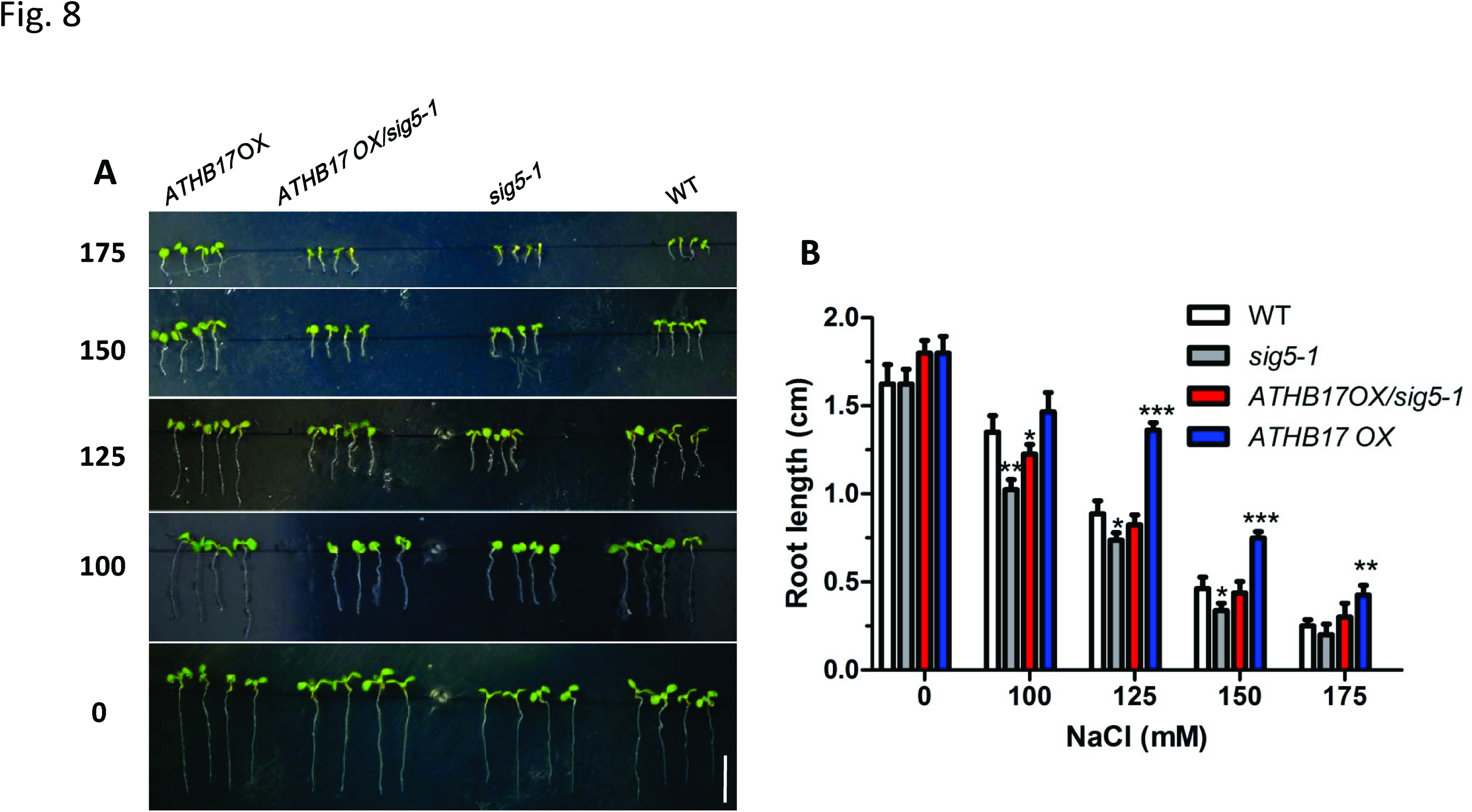
ATSIG5 acts downstream of ATHB17. (A) The phenotypes of *ATHB17 OX, ATHB17 OX/sig5-1, sig5-1* and WT seedlings grown on MS medium containing different concentrations of NaCl. Seeds were sowed on MS medium containing 0, 100, 125, 150 and 175 mM NaCl and grew vertically for 10 days. Bar = 1 cm. (B) Root length of the plants grown on MS medium containing different concentrations of NaCl.

Moreover, we analyzed the expression of the 20 PEGs, which were activated by ATHB17, in *sig5-1* and *ATSIG5* OX background. The results show that 8 of 20 PEGs (*PSBD*, *PSBC*, *PSAB*, *RBCL*, *PETB*, *RPOA*, *RPS8*, *YCF1.2*) had increased transcripts in *ATSIG5*-overexpressing plants while had decreased transcripts in the *sig5-1* mutant compared with WT. However, compared with WT, *PSBA*, *PSAA*, *PSBB* were found down-regulated and *RPS19*, *NDHD*, *NDHG* were found up-regulated both in *ATSIG5*-overexpressing and knockout plants, but with a much more dramatic change in the *sig5-1* mutant plants (Fig. 9). The expression change patterns of these PEGs partly agree with the expression pattern of these genes in *ATHB17* OX and KO plants, indicating that ATHB17-affected PEGs expression may be partly through ATSIG5.

**Figure 9.**
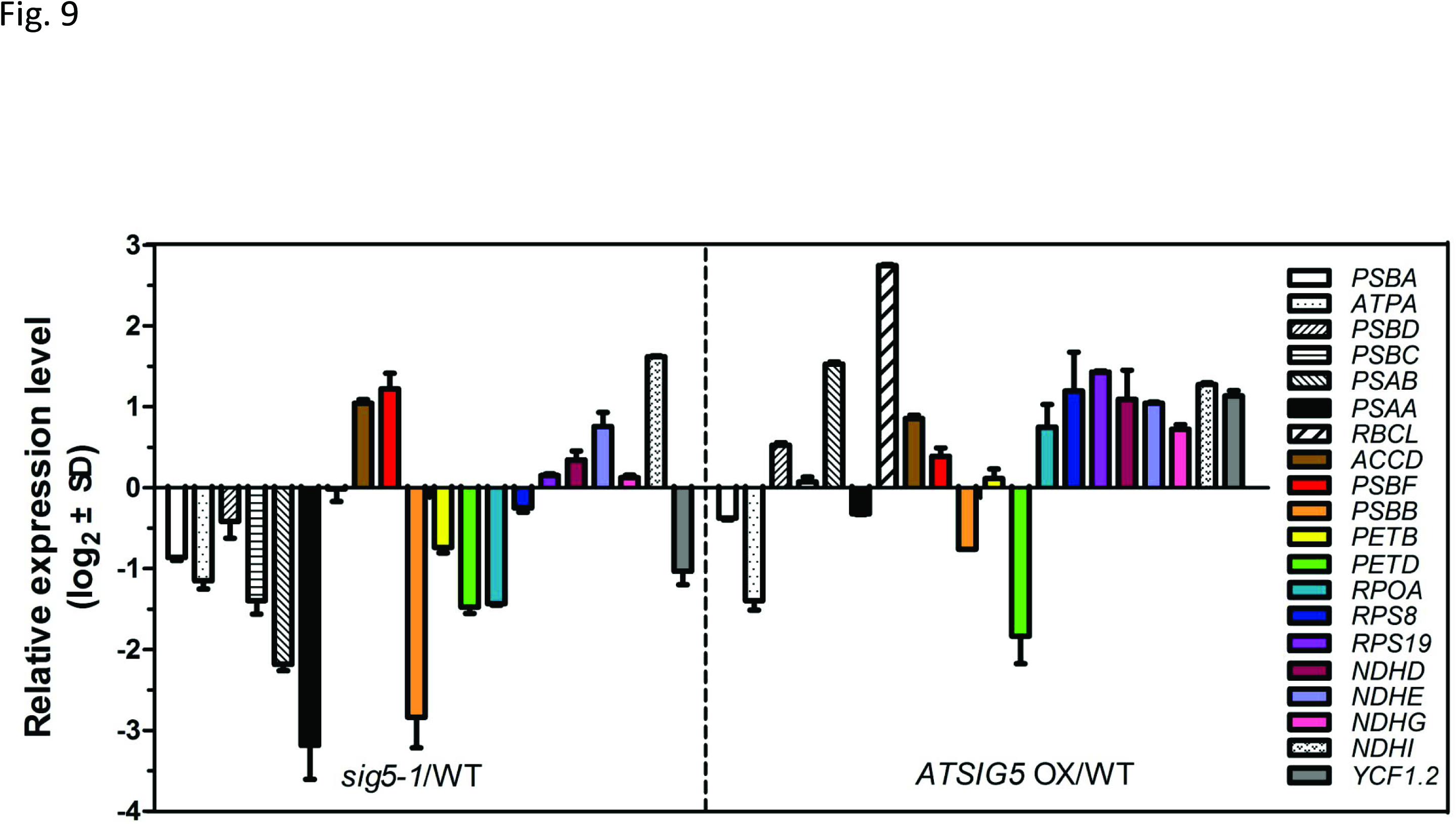
The expression levels of PEGs in *ATSIG5* OX and *sig5-1* mutant plants. 10-day-old plants grown under normal condition were used for qRT-PCR analysis. *UBQ5* was used as an internal control. Relative transcription levels of the genes in *ATSIG5 OX* and *sig5-1* plants are normalized to levels in the WT plants (WT = 0). Values are mean ± SD of three independent experiments.

## Discussion

Environmental stresses are great challenges for the development and growth of plants. In addition to developmental changes, chloroplasts constantly experience changing environment, thus a tight coordination between the nucleus and chloroplast is crucial to the survival of plant. These genome-coordinating mechanisms have been achieved through both anterograde (nucleus to organelle) and retrograde (organelle to nucleus) signals^43^. Most of chloroplast protein are nuclear-encoded, and the concentrations of these proteins are efficiently regulated by nuclear transcription^44^. TFs play important roles in the nuclear-chloroplast communication. Nuclear-encoded sigma factors regulate PEP activity to regulate the expression of different sets of genes responding to the external environmental signals^45^. Besides sigma factors, very few TFs have been isolated that regulate transcripts of nuclear photosynthetic genes and chloroplast genes. Up to date, GATA-type TFs GNC and CGA1 were reported to modulate the expression of chloroplast protein genes *GUN4* (*Genomes Uncoupled 4*) and *HEMA1* ^46^. ABI4 (Abscisic Acid Insensitive 4) represses the expression of photosynthetic nuclear genes, potentially acting as a master switch required for the modulation of nuclear genes in response to environmental signals, as well as developmental cues^47^. HYR (HIGHER YIELD RICE) is a master regulator in rice to environmental stress, directly activating several photosynthesis genes by binding to their promoters^48^. In addition, GLK1 and GLK2 TFs coordinate expression of the photosynthetic apparatus genes in *Arabidopsis* ^49^.

In this study, we found ATHB17, a HD-ZIP TF, played important roles in coordinating nuclear and chloroplast encoded photosynthetic gene expression in response to various abiotic stresses. By genetic analysis with knockout mutants and overexpression lines, we demonstrated that *ATHB17* was a positive regulator in response to abiotic stress. RNA-seq profiling analysis revealed that ATHB17 played as a repressor of NEPGs and an activator of PEGs. ATHB17 was reported as a transcriptional repressor containing EAR (ERF-associated amphiphilic repression)-like motif^40^. Consistent with this result, we found ATHB17 repressed the transcription of 26 NEPGs both under normal and salt stress conditions (Table 2 and Fig. 4), with 5 of them (*FDA6*, *LHCA2*, *LHB1B1*, *LHB1B2* and *PSBO1*) could be directly regulated by ATHB17 (Fig. 5). On the other hand, ATHB17 seems to activate the expression of many PEGs. Under normal condition, expression of 18 PEGs were found up-regulated in *ATHB17* OX while down-regulated in *ATHB17* KO compared WT. However, after salt stress, 13 PEGs still had higher expression in the *ATHB17* OX compared with WT and *ATHB17* KO plants, whereas only small expression difference of these genes existed between the WT and *ATHB17* KO plants (Table 3 and Fig. 4), indicting *ATHB17* regulating theses genes probably through an indirect way. Under salinity condition, other genes or the ATHB17 downstream signal pathway might be activated to recover the expression of these PEGs in ATHB17 KO plant. Interestingly, we found that ATHB17 could directly activate *ATSIG5* transcript (Fig. 6), which is an important nuclear encoded sigma factor modulating a number of PEGs, such as *PSBBT*, *PSBD*, *PSBC*, *PSBZ*, *PSAAB* and *PSBA* ^18,50-52^. Very likely, ATHB17 may regulate the expression levels of PEGs partially through *ATSIG5*, which is supported by genetic analysis (Fig. 8). Taken together, these studies demonstrate that ATHB17 acts as a regulator for coordinating the transcription of NEPGs and PEGs in plants under both normal and salt stress conditions.

Previous studies on salt stress tolerance mainly focused on genes related ion homeostasis, metabolites or osmoprotectants, antioxidant, hormones, ABA synthesis and signaling, and stress-responsive TFs^53,54^. However, so far very few researches focus on the coordinating photosynthetic genes to improve plant salt tolerance. Anyway, we did not found any difference in the expressed classic genes related to salt resistance between *ATHB17* OX and KO plants, including many stress-responsive genes, ABA synthesis and signaling pathway genes, as well as SOS genes (Table 1 and Fig. 3). These results partially agree with the results by Park et al.^41^, who reported that *ATHB17* overexpression did not affect the expression of a number of ABA-responsive genes. Thus, ATHB17-confered salt stress resistance may not through the traditional pathways.

Instead, we found that ATHB17 could coordinate the transcription of NEPGs and PEGs to deal with salinity stress. Firstly, ATHB17 represses the expression of genes related to light-harvesting complexes (*LHCA1, LHCA2*, *LHCA3, LHCA4*, *LHCA5, LHCB3*, *LHCB5*, *LHCB6*, *LHCB7, LHB1B1*, *LHB1B2*), chlorophyll a/b-binding proteins (*CAB1*, *CAB3*), PSIIoxygen evolving complex (*PSBO1*, *PSBO2*, *PPL1*, *PPL2*), thus reducing light harvest to alleviate photo-oxidative damage. Light-harvesting protein complex together with chlorophyll captures light energy and deliver it to the photosystems. Under stress condition, light harvest should be reduced, otherwise the photosystems would be overexcitated and damaged. Therefore, we can speculate that declining light capture by ATHB17 is one of the ways to protect plant from photodamage.

Secondly, ATHB17 is involved in the PS II repair cycle by regulating the genes encoding PS II subunits. The activity of PS II can be inactivated by a variety of environmental stresses, for these stresses inhibiting the repair of PS II rather than directly attacking it^55, 56^. D1 and D2, which are encoded by chloroplast gene *PSBA* and *PSBD*, bind all the redox-active components related to electron transfer of PS II and create oxidative power to break water molecules. Therefore, D1 and D2 are the main targets of oxidative damage^57, 58^. Thus, *PSBA* and *PSBD* need to express deviating from other chloroplast genes under various conditions, which might support the rapid turnover of D1 and D2 proteins in the PS s, which^59, 60^. From our study, we found that the expression of several genes encoding subunit of PS expression o*PSBD*, *PSBC* and *PSBB*, were increased in the *ATHB17* OX plants compared with WT and *ATHB17* KO plants under salt stress condition. However, the expression of *PSBA* in *ATHB17* OX is not significantly different from WT and *ATHB17* KO plants after salt stress treatment, while significantly up-regulated in *ATHB17* OX plants under normal condition. The high expression of *PSBD* and other PS II subunit genes may be beneficial for the efficient PS II repair under stress conditions.

Thirdly, ATHB17 enhanced the functions of a key antioxidant system in chloroplast to alleviate of salt-induced oxidative stress. Chloroplastic NAD(P)H dehydrogenase is involved in cyclic electron transport, chlororespiratory process, mitigating over reduction of the stroma, and play as key antioxidative enzymes to protect plants against oxidative damage under stress conditions^61-63^. A few of chloroplastic NAD(P)H dehydrogenase genes, including *NDHD*, *NDHI*, *NDHG*, were found had much higher expression levels in *ATHB17* OX plants compared with WT and *ATHB17* KO plants. Higher expression of these genes may help the *ATHB17* OX plants to alleviate the oxidative damage associated with salt stress.

At last, ATHB17 may be involved in balancing the stoichiometry of PS I to PS II under salt stress condition. By RNA-seq profiling analysis, we found that ATHB17 represses the expression of several nuclear encoded PS I subunits genes (*PSAF*, *PSAH1*, *PSAH2*, *PSAD1*), while activated the expression of chloroplast encoded PS II subunit genes (*PSBD*, *PSBC*, *PSBB*, *PSBA*, *PSBF*). It is well-known that PS II is the main target of oxidative damage under stress conditions, therefore, the subunit proteins of PS II are maintained with a rapid turnover rate to facilitate repair cycle^57,58^. Although the PS II in chloroplasts undergoes a frequent repair cycle, the functional PS II is decreased compared with that under normal condition. However, PS I maintains much higher stability compared with PS II under stress conditions. Thus, in order to balance the stoichiometry of photosystem I to photosystem II under salt stress condition, expression of genes encoding subunits of PS I should be simultaneously repressed to reduce the amount of PS I.

More interestingly, *ATHB17* is a multiple stress responsive gene, and can be induced by osmotic stress, PQ and ABA besides NaCl (Fig. 1C and D). Overexpression of *ATHB17* could also enhance plant tolerance to drought and oxidative stresses (Supplementary Fig. S4), indicating a similar mechanism as that of salt stress may exist in responding to these stresses. Taken together, our results imply that ATHB17 is an important TF regulating both NEPGs and PEGs to decline light harvest, enhance photosystem repair and antioxidative ability, and balance photosystem stoichiometry under stress conditions, thus improving the tolerance of plant to multiple stresses.

In *Arabidopsis*, the multiple-stress responsive plastid sigma factor ATSIG5, is structurally distinct from other sigma factors^64^. It is not only involved in the response of plant to blue light, but also regulates salt tolerance of plant by affecting repair of the PS II reaction center^18^. It may combine extrinsic and intrinsic signals important in adjusting plastid and nuclear gene expression upon light and environmental stress^50^. In this study, we found that *ATHB17* is also a multiple-stress responsive TF, similar with *ATSIG5*. NaCl, mannitol, PQ, ABA could highly induce its expression in leaves. Through Y1H and ChIP assay, we found ATHB17 could directly bind to *ATSIG5* promoter to regulate its transcription (Fig. 6). Overexpression of *ATSIG5* in *Arabidopsis* enhanced its salt tolerance, while *sig5* knockout plants became salt sensitive (Fig. 7). However, when *ATHB17* overexpressed in the *ATSIG5* knockout background, salt tolerance of the plants was partially impaired (Fig. 8). Moreover, the expression patterns of many PEGs in *ATSIG5* OX and *sig5* mutant compared with WT were similar to that of these genes in ATHB17 OX and KO compared with WT (Fig. 9). These results indicated that ATHB17 affects salt resistance partly depending on ATSIG5.

In conclusion, our study demonstrated that ATHB17 is an important TF in coordinating the expression of NEPGs and PEGs to cope with multiple stresses as illustrated in the working model (Fig. 10). Under stress condition, the expression of *ATHB17 is* induced, especially in leaves. As a TF, ATHB17 represses the transcription of many NEPGs indirectly or directly by binding to their promoters. Meanwhile, it directly activates transcription of *ATSIG5*, whose protein is then translocated to chloroplast as an important regulator of many chloroplast genes, such as *PSBA, PSBDC* and *PSBBT.* However, it may modulates several other chloroplast genes through other unknown pathways. Overall, it eventually reduces light harvest under stress, enhances photosystem repair cycle and antioxidative ability, and balances the stoichiometry of PS **I** to PS **II** in the chloroplast, therefore alleviating the damage to chloroplast under stress conditions and improving plant stress tolerance.

**Figure 10.**
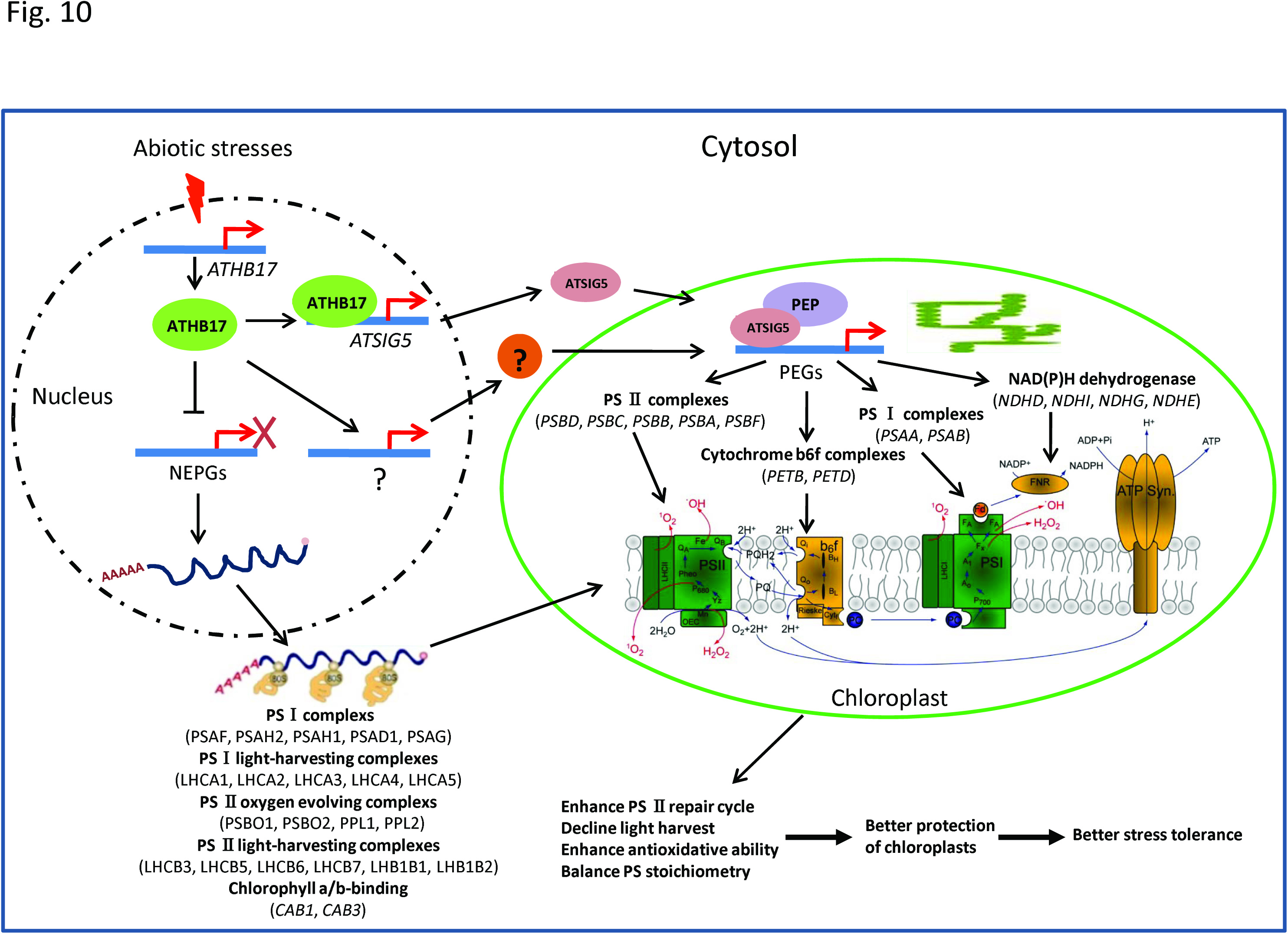
Proposed model: ATHB17 coordinates NEPGs and PEGs expression to cope with the abiotic stresses. Multiple abiotic stresses, such as salinity, drought and oxidative stress can significantly induce *ATHB17* expression. The ATHB17 protein can directly repress by bind to the promoters or indirectly repress the transcript of many NEPGs encoding for the subunits of photosystem complexes, which are then imported to the chloroplast. ATHB17 can also bind to the promoter of *ATSIG5 to* activate its transcript. ATSIG5 proteins are synthesized in the cytosol and subsequently targeted to the chloroplast, where it binds to PEP to tune the expression of several PEGs. Other unknown pathways modulated by ATHB17 may also exist to activate the expression of other PEGs. By coordinating the transcription of NEPGs and PEGs, ATHB17 enhance PS _ repair cycle and antioxidative ability, declines light harvest and balances PS stoichiometry, eventually reducing the damage to chloroplast and enhancing stress tolerance of the plant under stress conditions.

## Methods

### Plant materials and growth conditions

*Arabidopsis thaliana* (Col-0) and tobacco *(Nicotiana tobaccum*, NC89) were used for transformation. Seeds were sterilized in 10% bleach for 10 min, continued to wash five to six times with sterile water. For *Arabidopsis*, the seeds were first treated in 4 **°C** for 3 days vernalization, continued to sow on MS medium and 7-day-old *Arabidopsis* seedlings were transferred to soil. For tobacco, the washed seeds were directly germinated on MS medium, and 14-day-old seedlings were transferred to soil. The grown house condition was controlled at 22 ± 1 °C in a photoperiod of 16 h light and
8 h darkness.

### Constructs and preparation of transgenic plants

To get 35S::*ATHB17* and 35S::*ATSIG5* overexpression binary vector, the *ATHB17* or *ATSIG5* cDNA was isolated by RT-PCR with *ATHB17*-attb-LP and *ATHB17*-attb-RP or *ATSIG5*-attb-LP and *ATSIG5*-attb-RP and cloned into pCB2004^65^ via the GATEWAY cloning system. To analyze the *ATHB17* expression pattern, the 3.0 k bp promoter fragment was amplified with the primers p*ATHB17*-LP and p*ATHB17*-RP and then shuttled into the vector pCB308R^65^. For ChIP assay, *ATHB17* full-length coding sequence amplified by RT-PCR using specific primers *ATHB17*-HA-attb-LP and *ATHB17*-attb-RP was inserted into pCB2004 to get *pCB2004::ATHB17-HA* by the GATEWAY cloning system. All the constructs were electroporated into *Agrobacterium tumefaciens* C58C1, which were used to transform the wild type *Arabidopsis* plants as described^66, 67^. All the primers used are listed in Supplementary Table 1.

### Identification of *ATHB17* and *ATSIG5* knockout mutants and overexpression plants

A T-DNA insertion site of the SALK_095524 mutant in *ATHB17* ORF was confirmed by PCR using three specific primers: SALK_095524-LP, SALK_095524-RP, LBb1.3. Similarly, *sig5-* 1 (SALK_049021) mutant was identified by genomic PCR with three primers: *sig5-1* LP, *sig5-1* RP and LBb1.3. Mutant of *sig5-* 4 (SALK_101921) was identified by genomic PCR with three primers: *sig5-4* LP, *sig5-4* RP and LBb1.3. For expression analysis of the knockout mutants and overexpression transgenic plants, semi-RT-PCR was used with primers for the full-length coding sequence. All the primers used are listed in Supplementary Table 1.

### GUS activity staining assay

GUS staining of *pATHB17::GUS* transgenic plants was performed as described^68^. *Arabidopsis* seedlings were soaked in staining buffer containing 1 mM 5-bromo-4-chloro-3-indoryl-β-D-glucuronide (X-Gluc, Rose Scientific Inc., Somerset, NJ, U.S.A.) for overnight, then decolored using ethanol of different concentrations gradually. Individual representative seedlings were photographed.

### Hydroponic culture

20-day-old seedlings germinated on MS agar medium were used for hydroponic culture. The root of each tobacco seedling was wrapped in a sponge strip and inserted into the hole of floating body made by thick polystyrene foam board. The floating body floated on MS hydroponic solution with or without NaCl. Plants were cultured at 22 **°C** under 16-h light/8-h dark photoperiod. Nutrient solution was changed every 7 days.

### RNA-seq

Two-week-old seedlings of *ATHB17* OX, KO, and the WT control were treated with or without 200 mM NaCl in liquid MS medium, 60 rpm shaking in the air for 6 h, then all the six plant materials were harvested for total RNA extracting with TRIzol
reagent (Invitrogen Inc.). RNA-seq was performed and analyzed by BGI (Beijing
Genome Institute, Shenzhen) corporation following the protocol provided by the manufacturer. For annotation, all clean tags were mapped to the reference sequence and allowed no more than one nucleotide mismatch. Clean tags mapped to reference sequences from multiple genes were filtered. For gene expression analysis, the number of clean tags for each gene was calculated and then normalized to the number of transcripts per million tags (TPM).

### qRT-PCR

Total RNA was extracted using TRIzol reagent (Invitrogen Inc.), first-strand cDNA was synthesized from 1 µg total RNA in a 20 µl reaction mixture with Prime Script RT regent kit (TAKARA BIOTECHNOLOGY CO., LTD). The transcript levels of classic stress-related genes and other genes were examined with specific primers. The PCR was performed on ABI step-one instrument with the amplification conditions for 40 cycles of 95 °C for 1 min, 62 °C for 20 seconds and 72 °C for 30 seconds. *UBQ5* was used as the internal control. The relative expression levels were calculated by the 2^-ΔΔCT^ method. All the primers used are listed in Supplementary Table 1.

### Yeast one-hybrid assay (Y1H)

A cDNA fragment encoding *ATHB17* was amplified by the primers: the forward with restriction endonuclease BamH I, 5’-<u>CGGGATCC</u>GACCAGCTAAGGC TAGACATGAA-3’and the reverse with restriction endonuclease Xba I, 5’-<u>GCTCTAGATCA</u>ACGATCACGCTCTTGCG-3’, and inserted into plasmid pAD-GAL4-2.1 (pAD) to get AD/ATHB17Δ107.

To get the report vectors containing HD-binding sequences, three copies of the HD-binding sequence, containing Sac I and Mlu I adaptors, were annealed and inserted into the Sac I and Mlu I sites of pHIS2. To get the report vectors containing promoter sequence of different genes, 25 bp promoter segment containing HD-binding sequence with Sac I and Mlu I adaptors was annealed and inserted into pHIS2, respectively. The constructs were confirmed by sequencing. The pAD and pHIS2 empty vector were used for negative control. AD/ATHB17Δ107 and the reporter pHIS2 containing different DNA sequence were co-transfected into yeast cells Y187 respectively. The yeast was first grown on SD-Trp-Leu medium for 3 days at 30°C,. and then transferred to SD-Trp-Leu-His medium with 10 mM or 20 mM 3-aminotriazole (3-AT, sigma) at different dilutions. The yeasts were incubated at 30 °C for 5 days and the extent of yeast growth was determined.

### ChIP-PCR assay

Leaves of 10-day-old T_3_ homozygote of *35S::ATHB17-HA* transgenic plants and anti-HA tag antibody (Cali-Bio, CB100005M) were used for pulling down the chromatin, mainly as previously described^69^. After degrading the associated proteins with proteinase K, the chromatin DNA samples were treated using phenol/chloroform, then precipitated and finally eluted in 30 μl TE buffer. ChIP-PCR was then used to verify the promoter segment of related genes using the primers listed in Supplementary Table 1.

### EMSA assay

ATHB17Δ107 protein was expressed in *Escherichia coli* with pET28a (+) protein expression system (Novagen) and purified with Ni^2+^ chromatography. Three copes of complementary single-stranded HD-binding sequence were synthesized and annealed to form double-stranded DNA fragment. The DNA fragments were marked with α-^32^P-dCTP and gel purified as probes. Purified ATHB17Δ107 protein (200 ng) was incubated with probes for 30 min on ice. For the competition test, non-labeled probe and non-specific probes were added into the binding reaction. Each reaction was loaded on a 4.5% native polyacrylamide gel with 0.5 × TBE buffer. The gel then exposed to X-ray film.

### Statistical analysis

Statistically significant differences (P < 0.05 or P < 0.01 or P < 0.001) were computed based on the Student’s *t*-tests.

## Acknowledgements

This work was supported by MOST (Grant # 2012CB114304 to C.-B.X) and NNSFC (Grant # 30700051 to C.-B.X). We thank ABRC for offering T-DNA insertion lines.

## Author contributions

P.Z., R.C. and L.-H.Y. performed the experiments and data analysis. P.X. and Y.C. contributed to RNA-seq data analysis. J.-L.M. assisted with the experiment. L.-H.Y. and P.Z. wrote the manuscript. C.-B.X. and L.-H.Y. conceived the study and designed the experiments. C.-Z.Z. contributed to ATHB17 protein purification and EMSA experiment. C.-B.X. supervised the project and edited the manuscript.

## Additional information

Supplementary Fig. S1. Identification of *ATHB17* OX and KO.

Supplementary Fig. S2. Functional complementation assay.

Supplementary Fig. S3. Overexpression of *ATHB17* improved salt tolerance of tobacco plants.

Supplementary Fig. S4. ATHB17 is involved in drought and oxidative stresses.

Supplementary Fig. S5. ATHB17 binding to the HD-binding cis-elements.

Supplementary Fig. S6. Identification of *ATSIG5* overexpression lines and knockout lines.

Supplementary Table 1. Primer sequences used in this study.

Supplementary Data 1. Gene Ontology term enrichment analysis of 426 down-regulated genes of *ATHB17* OX/*ATHB17* KO under NaCl treated condition.

Supplementary Data 2. Gene Ontology term enrichment analysis of 1704 down-regulated genes of *ATHB17* OX/*ATHB17* KO under normal condition.

